# Glacial runoff promotes deep burial of sulfur cycling-associated microorganisms in marine sediments

**DOI:** 10.1101/661207

**Authors:** Claus Pelikan, Marion Jaussi, Kenneth Wasmund, Marit-Solveig Seidenkrantz, Christof Pearce, Zou Zou Anna Kuzyk, Craig W. Herbold, Hans Røy, Kasper Urup Kjeldsen, Alexander Loy

**Affiliations:** Division of Microbial Ecology, Centre for Microbiology and Environmental Systems Science, University of Vienna, Vienna, Austria; Austrian Polar Research Institute, Vienna, Austria; Center for Geomicrobiology and Section for Microbiology, Department of Bioscience, Aarhus University, Aarhus, Denmark; Department of Geoscience, iClimate, and Arctic Research Centre, Aarhus University, Denmark; Department of Geological Sciences, University of Manitoba, Winnipeg, Canada

**Keywords:** sulfate-reducing microorganisms, marine sediment, glacial impact, deep biosphere, microbial community assembly, Greenland, arctic

## Abstract

Marine fjords with active glacier outlets are hot spots for organic matter burial in the sediments and subsequent microbial mineralization, and will be increasingly important as climate warming causes more rapid glacial melt. Here, we investigated controls on microbial community assembly in sub-arctic glacier-influenced (GI) and non-glacier-influenced (NGI) marine sediments in the Godthåbsfjord region, south-western Greenland. We used a correlative approach integrating 16S rRNA gene and dissimilatory sulfite reductase (*dsrB*) amplicon sequence data over six meters of depth with biogeochemistry, sulfur-cycling activities, and sediment ages. GI sediments were characterized by comparably high sedimentation rates and had ‘young’ sediment ages of <500 years even at 6 m sediment depth. In contrast, NGI stations reached ages of approximately 10,000 years at these depths. Sediment age-depth relationships, sulfate reduction rates, and C/N ratios were strongly correlated with differences in microbial community composition between GI and NGI sediments, indicating that age and diagenetic state were key drivers of microbial community assembly in subsurface sediments. Similar bacterial and archaeal communities were present in the surface sediments of all stations, whereas only in GI sediments were many surface taxa also abundant through the whole sediment core. The relative abundance of these taxa, including diverse *Desulfobacteraceae* members, correlated positively with sulfate reduction rates, indicating their active contributions to sulfur-cycling processes. In contrast, other surface community members, such as *Desulfatiglans, Atribacteria* and *Chloroflexi*, survived the slow sediment burial at NGI stations and dominated in the deepest sediment layers. These taxa are typical for the energy-limited marine deep biosphere and their relative abundances correlated positively with sediment age. In conclusion, our data suggests that high rates of sediment accumulation caused by glacier runoff and associated changes in biogeochemistry, promote persistence of sulfur-cycling activity and burial of a larger fraction of the surface microbial community into the deep subsurface.

**Contribution to the Field Statement:** In most coastal marine sediments organic matter turnover and total energy flux are highest at the surface and decrease significantly with increasing sediment depth, causing depth-dependent changes in the microbial community composition. Glacial runoff in arctic and subarctic fjords alters the composition of the microbial community at the surface, mainly due to different availabilities of organic matter and metals. Here we show that glacial runoff also modifies microbial community assembly with sediment depth. Sediment age was a key driver of microbial community composition in six-meter-long marine sediment cores from the Godthåbsfjord region, south-western Greenland. High sedimentation rates at glacier-influenced sediment stations enabled a complex community of sulfur-cycling-associated microorganisms to continuously thrive at high relative abundances from the surface into the sediment subsurface. These communities consisted of putative fermenters, sulfate reducers and sulfur oxidizers, which likely depended on high metal concentrations in the relatively young, glacier-influenced sediments. In non-glacier-influenced sediments with lower sedimentation rates, these sulfur-cycling-associated microorganisms were only present near the surface. With increasing sediment depth these surface microorganisms were largely replaced by other surface microorganisms that positively correlated with sediment age and belong to known taxa of the energy-limited, marine deep biosphere.

## Introduction

Arctic fjords with marine-terminating glaciers constitute an important interface for freshwater and sediment influx from land into the sea, thereby influencing the physical and chemical conditions in the coastal marine ecosystems (Etherington et al., 2007; Svendsen et al., 2002). The high influx of sedimentary materials, e.g., minerals, terrigenous organic matter and metals, in glacier-associated fjords has a strong effect on the distributions of the benthic microbial communities (Bourgeois et al., 2016; Buongiorno et al., 2019; Park et al., 2011). Increased water turbidity in close proximity to the glacier can negatively influence surface water primary production (Etherington et al., 2007; Zajączkowski, 2008), which leads to lower organic matter availability in the underlying sediments (Bourgeois et al., 2016). High sediment accumulation rates often seen in such glacier-proximal environments are also limiting benthic life. On the other hand, glacial meltwater also provides an important source of dissolved nutrients, which can stimulate phytoplankton growth beyond the high turbidity zone (Meire et al., 2015; Sørensen et al., 2015; Statham et al., 2008). Consequences of increased primary production together with strong sediment supply are a net CO_2_ uptake in glaciated fjords as well as rapid burial of fresh detrital phytoplankton biomass to the underlying sediments (Bourgeois et al., 2016; Meire et al., 2015; Smith et al., 2015). Fjord sediments also receive significant amounts of terrigenous organic matter as evidenced by high C/N ratios of the sediment organic matter pool (Goñi et al., 2013; Wehrmann et al., 2014). With these large inputs of both marine and terrigenous organic matter, the ultimate role of glaciated fjord sediments in the global carbon cycle depends on the extent to which the large organic matter inputs get degraded, and thus there is a need to better understand constraints on microbial community structure and degradation potential.

The labile fraction of organic matter in marine sediments is initially degraded by microorganisms with hydrolytic and fermenting capabilities (Müller et al., 2018). These largely unidentified microbial species typically excrete a wide range of enzymes, which enable rapid organic matter turnover even in cold arctic sediments (Arnosti et al., 1998; Teske et al., 2011). Fermentation products released by primary organic matter degrading microorganisms are either further degraded by secondary fermenters or are mineralized completely to CO_2_ via microbial respiration (Arndt et al., 2013).

Of prime importance in microbial organic matter degradation is the respiratory reduction of sulfate to sulfide, which facilitates up to 69% of the total organic matter mineralization in Arctic fjord sediments (Sørensen et al., 2015). Other key electron acceptors are metals such as iron(oxyhydr)oxides and manganese oxides that can be introduced in high amounts via glacial runoff to marine sediments and are subjected to redox cycling (Bhatia et al., 2013; Buongiorno et al., 2019; Laufer et al., 2016; Wehrmann et al., 2014). Fe(III) and Mn(IV) also facilitate the oxidation of reduced sulfur compounds (Wehrmann et al., 2014; Zopfi et al., 2004) and thereby fuel a cryptic sulfur cycle in glacier-influenced sediments (Wehrmann et al., 2017).

In sediments with comparably slow sediment accumulation, which typify non-glaciated Arctic shelf areas (Kuzyk et al., 2013), the rate of organic matter mineralization is highest at the surface and decreases significantly with increasing sediment depth (Kuzyk et al., 2017; Lomstein et al., 2012), reflecting a concomitant decrease in energy available for cell maintenance and growth (Starnawski et al., 2017). This depth gradient results in pronounced compositional changes in the benthic microbial community, which are driven by highly selective survival of microorganisms that are able to subsist in the energy-limited subsurface (Bird et al., 2019; Marshall et al., 2019; Petro et al., 2017; Starnawski et al., 2017). Here, we hypothesized that this strong environmental filtering effect would be attenuated in sediments with high rates of sedimentation (i.e., in coastal sediments impacted by glacial runoff) and result in a different pattern of microbial community assembly with depth. Therefore, we investigated microbial diversity and community compositions of glacier-influenced (GI) and non-glacier-influenced (NGI) coastal sediments in the Godthåbsfjord region of Greenland. Compositions of bacterial and archaeal communities and the community of putative sulfite/sulfate-reducing microorganisms (SRM), as analyzed by 16S rRNA and dissimilatory sulfite reductase (*dsrB*) gene amplicon sequencing, respectively, were compared across sediment samples of different depths and ages. Co-occurrence analyses of operational taxonomic units (OTUs) and correlations with biogeochemical data revealed key environmental factors that were driving the major community differences between GI and NGI sediments. These analyses also identified various uncultivated microorganisms that were associated with sulfur cycling. As hypothesized, our results demonstrate that glacial runoff exerts a strong influence on microbial community assembly processes and community functions in marine sediments.

## Materials and Methods

### Sediment sampling

The sediment cores used in this study were collected using a gravity corer in 2013 during a research cruise on board of RV *Sanna* (Seidenkrantz et al., 2014). Sampling of the cores for this study was described previously (Glombitza et al., 2015). In brief, up to 6 m long gravity cores were recovered from four sites on the open shelf and within the Godthåbsfjord (Nuup Kangerlua) system in South West Greenland (Table 1, Supplementary Figure S1) in August 2013. The four stations can be broadly subdivided into two groups. First, the non-glacier-influenced (NGI) stations 3 and 6. Station 3 (core SA13-ST3-20G) is located outside the fjord on the continental shelf of the Labrador Sea. Station 6 (core SA13-ST6-40G) is situated in the Kapisigdit Kanderdluat, a side-fjord without glaciers. Second, the glacier-influenced (GI) stations 5 and 8. Station 5 (core SA13-ST5-30G) is located in the main channel of the Godthåbsfjord. Although station 5 is not directly in front of a glacier, most glacier-derived material is transported towards the Labrador Sea across this site. Station 8 (core SA13-ST8-47G) is in very close proximity to a glacier front at the northernmost outlet of the Greenland ice sheet in the Kangersuneq fjord (Supplementary Figure S1).

**Table 1.** Description of the sampling stations and cores, including sampling position, water depth and core length. Table was modified from Glombitza et al., 2015. The exact location of the sampling stations is indicated in Supplementary Figure S1.

| Station | Core name | Latitude (N) | Longitude (W) | Waterdepth (m) | Core length (cm) |
| --- | --- | --- | --- | --- | --- |
| 3 | SA13-ST3-20G | 6426.7425' | 5247.6486' | 498.2 | 587 |
| 5 | A13-ST5-30G | 6425.3479' | 5130.6209' | 622.4 | 607 |
| 6 | SA13-ST6-40G | 6429.0604' | 5042.3240' | 389 | 562 |
| 8 | SA13-ST8-47G | 6440.7078' | 5017.4672' | 475.8 | 569 |

For molecular analyses, sediment cores were sampled by cutting small holes into the core liner and scraping off the surface with sterile spatulas before collecting sediment samples in duplicates with sterile 5 mL cut-off syringes. The sediment sub-samples were packed in Whirl-Pak bags and immediately frozen at −80°C. Porewater was extracted from the sediment cores as described previously (Glombitza et al., 2015).

### Analytical methods

Sulfate, hydrogen sulfide, and dissolved inorganic carbon (DIC) concentration measurements, as well as sulfate reduction rate (SRR) measurements were previously described in Glombitza et al. (2015). In addition, ferrous iron, total organic carbon (TOC), and total nitrogen (TN) were analyzed. For analyses of ferrous iron (Fe^2+^) concentrations, porewater was amended with 0.5 M HCl (1:1, v/v) and stored at 4°C. One mL of ferrozine solution (1 g L^-1^ in 50 mM HEPES-buffer at pH 7) was subsequently added to 20 μL of acid-preserved porewater, producing a magenta color reaction (Phillips and Lovley, 1987; Sørensen, 1982). The absorbance was measured at 562 nm (Stookey, 1970) with a spectrophotometer (FLUOstar Omega, BMG Labtech GMBH). To determine total organic carbon (TOC) and total nitrogen (TN), sediment samples were treated with sulfurous acid (5-6% w/w) to remove inorganic carbon. Once dried, 50 mg of acidified sediment were packed into cleaned tin cups and burned in the elemental analyzer (FLASH EA 1112 series, Thermo Scientific). TOC and TN concentrations were calculated from a standard curve with wheat flour, which contains 43.37% of carbon and 2.31% of nitrogen. The C/N ratio was calculated as the molar ratio of total organic carbon to total nitrogen. For determination of methane concentrations, 2 cm^3^ sediment was transferred to 20 mL GC vials containing 2.5 mL saturated NaCl with excess crystalline salt. The bottles were vigorously shaken to release methane into the headspace of the GC vials and stored upside down at −20°C until further analysis. Methane concentrations in the vial headspace were subsequently measured by gas chromatography (SRI 310C GC, SRI Instruments Europe GmbH) with a flame ionization detector. Ammonium concentrations were determined from 2 mL of porewater. Concentrations were analyzed spectrophotometrically as previously described (Dansk-Standard, 1975) (Bower and Holm-Hansen, 1980).

Sediment age profiles for cores from stations 3, 5, and 6 were based on radiocarbon dating and age modeling. Materials and depths were chosen for dating based on availability. For the ^14^C age determination, calcareous mollusk shells, benthic foraminifera, and seaweed samples were collected from cores and the ^14^C concentrations were determined by Accelerator Mass Spectrometry at the AMS ^14^C Dating Centre of Aarhus University. The ^14^C ages were calibrated using the Marine13 radiocarbon calibration curve (Reimer et al., 2013) with a reservoir correction of ΔR = 140 ± 35 years. Age-depth models for cores from station 3 and 6 were calculated using the software Oxcal v4.2 (Ramsey, 2008). The presence of rapidly deposited turbidite sequences in the core from station 5 made a detailed age model difficult.

In the core retrieved from station 8 no material usable for ^14^C dating was found, and thus radiocarbon dating was not possible. Instead, the sediment age was estimated from ^210^Pb and ^226^Ra measurements of freeze-dried, homogenized sediment samples. The water content of each sample was determined before and after freeze-drying, and results were reported on a dry-weight basis and salt-corrected for bottom-water salinity at the time of sampling. Core sections were analyzed by gamma spectroscopy using a CANBERRA^®^ Broad Energy Germanium with a P-type detector (model BE3830). Detector efficiency and self-absorption were corrected by counting reference material from the International Atomic Energy Agency (IAEA) within the same geometry. The reproducibility errors, determined by counting the same sample four times, were 5.7 and 3.8%, for ^210^Pb and ^226^Ra, respectively. ^210^Pb was also determined from its ^210^Po daughter isotope using alpha spectroscopy (n=23 sediment sections). The samples were prepared as previously described (Flynn, 1968). A ^209^Po tracer, calibrated against a ^210^Po NIST standard (Isotope Product Laboratories) was employed for quantitation. The reproducibility error was less than 1%. Sedimentation rates that were used for the age model presented in this study were estimated from the least-squares fit to the natural log of the excess ^210^Pb (^210^Pb*ex*) in the core and the output of a one-dimensional two-layer advection diffusion model that accounts for both bioturbation and compaction with depth (Kuzyk et al., 2015; Lavelle et al., 1985). When using any tracer data for age model reconstructions within marine sediments, it is important to recognize that profiles may be affected by mixing (physical or bioturbation). Physical disturbance such as a rapid deposition event (turbidite) may be seen in, for example, reversal in the sediment porosity gradient and unusually low ^210^Pb profiles in a particular layer. Many organic-rich shelf sediment cores exhibit two-layer ^210^Pb profiles reflecting a ‘surface mixed layer’ (SML) overlying accumulating sediments that are subject to little or no mixing (Kuzyk et al., 2013). If mixing rates are significant relative to sediment accumulation, then it is not possible to assign ages to specific sediment sections because each section will contain a distribution of various ages. In the case of station 8, the sediments are low in organics and ^210^Pbex decreases exponentially with depth, implying that mixing is minor relative to sediment accumulation. Furthermore, the sediment is highly laminated, indicating very little mixing.

To estimate the loss of surface sediment by gravity coring and to correct the age-depth model, we compared the porewater profiles of ammonium, dissolved inorganic carbon and the carbon isotope ratio ^13^C/^12^C of DIC (data not shown) between Rumohr cores collected during the same cruise and the gravity cores presented in this study. The upper 18 cm of the gravity core retrieved at station 3 and the upper 10 cm of the core retrieved at station 6 were missing. The depths of potential surface sediment loss at the two GI stations were corrected as the mean values of the two first cores (14 cm) since the Rumohr core casts at the GI stations were not successful. Finally, sediment age was recalculated as “actual age”, i.e., the surface of the seafloor was considered as 0 year old at the time of sampling, based on the age models and the above mentioned depth correction.

### DNA extraction and preparation of 16S rRNA gene and dsrB amplicon libraries

Approximately 0.5 to 1 g of sediment was used for DNA extraction according to a previously established protocol (Kjeldsen et al., 2007). Per station, 8 to 10 samples from different depths, corresponding to approximately 2 samples per meter core were selected for further analyses, with highest resolution at the top of the cores (Supplementary Table S1). Barcoded 16S rRNA gene amplicons were produced with a two-step PCR barcoding approach (Herbold et al., 2015), using the general bacterial and archaeal primers U519F (5’-CAGCMGCCGCGGTAATWC-3’) and 802R (5’-TACNVGGGTATCTAATCC-3’) for initial amplification (Klindworth et al., 2013). These primers were modified with a 16 bp head sequence as described previously (Herbold et al., 2015). The first round of amplification was performed in triplicates with 12.5 μL per reaction volume. The reaction mix contained 1× Taq buffer (Thermo Scientific), 0.2 mM dNTP mix (Thermo Scientific), 2 mM MgCl_2_, 0.25 U Taq polymerase (Thermo Scientific), 0.2 μM of each primer and approximately 1-10 ng DNA. The PCR started with a denaturation at 95°C for 3 min, followed by 30 cycles of 95°C for 30 s, 48°C for 30 s and 72°C for 30 s, and a final elongation at 72°C for 2 min. The subsequent barcoding PCR round (50 μL total volume) was performed with 1× Taq buffer (Thermo Scientific), 0.2 mM dNTP mix (Thermo Scientific), 2 mM MgCl_2_, 1 U Taq polymerase (Thermo Scientific), 0.2 μM of each primer and 2 μL of the pooled triplicate PCR products from the first PCR reaction. The thermal cycling program consisted of an initial denaturation at 95°C for 3 min, 12 cycles of 95°C for 30 s, 52°C for 30 s and 72°C for 30 s, followed by a final elongation at 72°C for 2 min. The *dsrB* amplicons were produced according to an established protocol (Pelikan et al., 2016). Barcoded amplicons were mixed and further prepared for multiplexed, paired-end MiSeq sequencing (Herbold et al., 2015). Sequence datasets are available in the NCBI Sequence Read Archive under study accession number PRJNA546002.

### Sequence data processing

16S rRNA gene and *dsrB* amplicon raw reads were demultiplexed, filtered and clustered as described previously (Herbold et al., 2015; Pelikan et al., 2016) using fastq-join (Aronesty, 2013) to merge reads and UPARSE version 8.1.1861 (Edgar, 2013) to generate OTUs. Phylum/class-level classification of 16S rRNA-OTUs was performed with the Ribosomal Database Project naïve Bayesian classifier in MOTHUR (Schloss et al., 2009; Wang et al., 2007), using the SILVA database v.128 (Quast et al., 2013) as a reference. *dsrB*-OTUs were classified by phylogenetic placement of representative sequences into a DsrAB reference tree (Müller et al., 2015) that was updated with novel sequences from diverse candidate phyla (Anantharaman et al., 2017; Hausmann et al., 2018; Parks et al., 2017). This DsrAB reference tree was constructed by de-replicating novel DsrA and DsrB sequences with less than 100 % similarity to any DsrAB sequence in the original reference database and aligning them to the reference alignments of DsrA and DsrB (Müller et al., 2015) using MAFFT (Katoh et al., 2002). The combined DsrA and DsrB alignments were then concatenated and sequences with a total length of less than 500 amino acids were removed. The concatenated DsrAB alignment was clustered at 70 % sequence identity with usearch (Edgar, 2010), and alignment positions were kept if they were conserved in at least 10 % of all sequences in the 70%-clustered alignment (56 sequences). The unclustered DsrAB alignment (2985 sequences) was then filtered for conserved alignment positions using seqmagick (https://fhcrc.github.io/seqmagick/) and was used to generate a maximum likelihood tree with FastTree (Price et al., 2010). 16S rRNA gene and *dsrB* OTU tables were processed in R using native functions (R Core Team, 2015) and the R software package phyloseq (McMurdie and Holmes, 2013). OTU counts were rarefied, i.e., sub-sampled at the smallest library size (*dsrB*: 1521; 16S rRNA gene: 4517) and transformed into relative abundances for all further analyses, except for network analyses, which were performed with the unrarefied OTU count matrices (Friedman and Alm, 2012).

### Statistical analyses

Shannon alpha diversity was calculated using R (R Core Team, 2015). Beta diversity analyses were performed with the R software package vegan (Oksanen et al., 2017), including calculations of Bray-Curtis distances with the function ‘vegdist()’ and nonmetric multidimensional scaling ordination analysis with the function ‘nMDS()’. Environmental variables were tested for effects on the overall community composition by Mantel tests using the native R function ‘mantel()’. Obtained p-values were corrected for multiple testing with the native R function ‘p-adjust()’ using the Benjamini-Hochberg correction method. Correlations of individual OTUs with environmental variables were calculated with the native R function cor() using the Spearman correlation coefficient. P-values were generated by permuting the values of each environmental variable followed by correlation of individual OTU abundances with the permuted environmental variable. This process was repeated 1000 times and the obtained p-values were corrected as described above.

Correlation network analyses were performed separately for 16S rRNA- and *dsrB*-OTUs to highlight potential synergistic interactions between microbial community members (Weiss et al., 2016). Species co-occurrence networks were calculated using SparCC (Friedman and Alm, 2012) based on count matrices of all OTUs with >10 reads in at least 7 out of 34 samples. Use of more abundant and prevalent OTUs increases sensitivity of the network analyses (Berry and Widder, 2014). P-values were generated as described above. Only positive OTU correlations >0.5 were considered. Attributes of individual OTUs, i.e., sampling station and sediment age at which the OTU was found at the highest relative abundance, were assigned to OTUs in R and networks were visualized in Cytoscape (Shannon et al., 2003). Significant OTU clusters, i.e., significantly more interactions between OTUs within the community cluster than with OTUs outside the community cluster, were defined by Mann-Whitney U tests using the Cytoscape plugin “clusterONE” (Nepusz et al., 2012).

### Phylogenetic analysis

Representative sequences of 16S rRNA-OTUs were aligned with the SINA aligner (Pruesse et al., 2012) using the SILVA database v.128 (Quast et al., 2013) as a reference. Sequences that were closely related to 16S rRNA-OTUs were extracted from the SILVA database and used to construct a reference tree with FastTree (Price et al., 2010). Subsequently, 16S rRNA-OTU sequences were placed into the reference tree using the EPA algorithm (Berger et al., 2011) in RAxML (Stamatakis, 2014). The placement trees of 16S rRNA-OTUs and *dsrB*-OTUs (utilized for *dsrB*-OTU classification) were visualized with iTOL (Letunic and Bork, 2007).

## Results

### Depth profiles of sediment age and porewater chemistry differ substantially between non-glacier-influenced and glacier-influenced sediments

A goal of the present study was to identify environmental factors (biogeochemical data is partially described in Glombitza *et al*., 2015) that shape the microbial community compositions and interactions in NGI and GI sediments. NGI stations 3 and 6 are located on the open shelf and within the Godthåbsfjord, respectively, and were both characterized by a strong gradient of sediment age due to comparably low sedimentation rates with maximum ages of the gravity cores close to 10,000 years (Figure 1). Furthermore, NGI sediments had high TOC and TN concentrations, as well as low C/N ratios down to 500 cm sediment depth. SRR decreased with depth at both stations and sulfate became depleted in the bottom of the core from station 3, but not in the core from station 6. At station 3, hydrogen sulfide concentrations gradually increased with depth and decreased again below a depth of 400 cmbsf, coinciding with the appearance of methane in the porewater. At station 6, hydrogen sulfide was present in lower amounts and methane did not accumulate at any depth.

**Figure 1.**
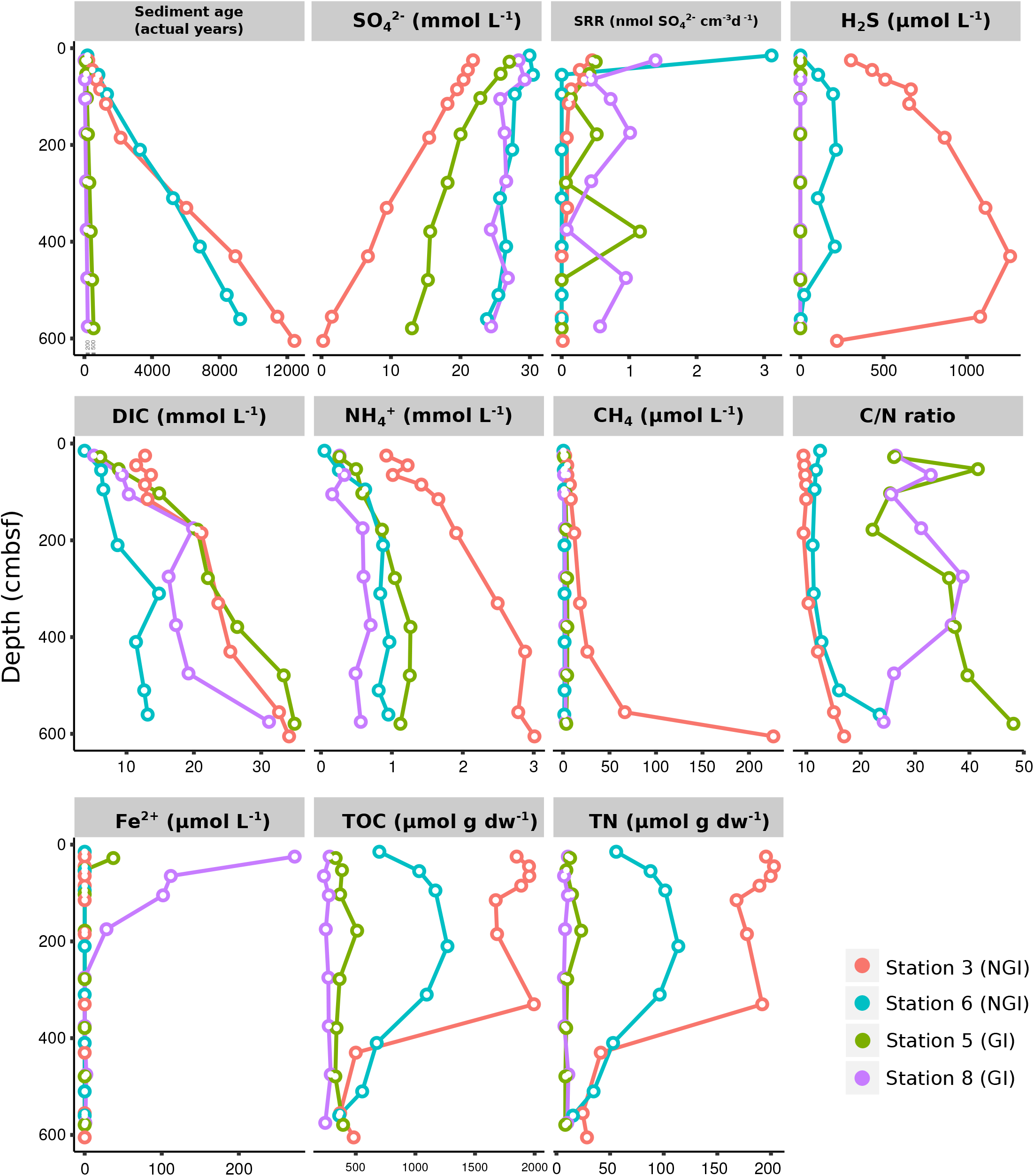
Physicochemical sediment properties in non-glacier-influenced (NGI) and glacier-influenced (GI) sediment cores. Colours indicate the sampling station. Note that the scales are different for each physicochemical parameter. Data on SRR and DIC, as well as, sulfate and sulfide concentrations were taken from Glombitza et al., 2015.

In comparison, GI stations 5 and 8 were both characterized by high sedimentation rates as indicated by the low sediment ages, i.e., around 200 years at the bottom of the core at station 8 and around 500 years at the bottom of the core at station 5 (Figure 1). GI stations had low TOC and TN concentrations, as well as high C/N ratios of up to 48. The porewater contained dissolved Fe^2+^ in the upper 250 cmbsf at both GI stations. SRR were high in deeper sediment layers at the GI stations. Notably, SRR in the deep sediments of station 8 were higher than those measured in the uppermost sediment samples from the core retrieved at station 3. Despite substantial SRR, hydrogen sulfide was not detected at any depth.

### Glacier runoff and sediment age are strong determinants of microbial community composition

In total, 6755 16S rRNA-OTUs and 1094 *dsrB*-OTUs were obtained by amplicon sequencing. NGI and GI sediments clearly differed in 16S rRNA- and *dsrB*-OTU compositions and these differences increased with depth at NGI stations (Supplementary Figure S2 A and B). Mantel correlations with environmental factors revealed that the 16S rRNA- and *dsrB*-OTU compositions were mostly impacted by sediment age and C/N ratio of organic matter (Table 2). Further differences in 16S rRNA gene and *dsrB*-OTU compositions were explained to a lesser extent by sediment structure (i.e., density and porosity), TOC, TN, SRR and Fe^2+^ concentrations (the latter two parameters were only significant for the 16S rRNA gene community) (Table 2). Gradually increasing sediment age with depth at NGI stations (Figure 1) was associated with gradual changes in 16S rRNA gene and *dsrB* beta-diversity with depth (Supplementary Figure S2 A and B). 16S rRNA and *dsrB* alpha-diversity at the NGI stations gradually decreased with depth (Supplementary Figure S2 C and D). In contrast, alpha-diversity in GI stations remained rather high throughout the cores (Supplementary Figure S2 C and D). Relative abundance patterns of most phyla/classes and DsrAB families at GI stations 5 and 8 did not follow a gradual change in compositions with increasing sediment depth like at NGI stations 3 and 6, but remained rather constant, with some fluctuations among taxa (Figure 2). *Alpha-, Delta-* and *Gammaproteobacteria, Campylobacterota* and notably *Cyanobacteria* were overall more abundant at GI sediments as compared to NGI sediments (Figure 2 A). Furthermore, the *dsrAB*-containing community in GI sediments had higher relative abundances of *Desulfobacteraceae*, *Desulfobulbaceae*, uncultured DsrAB family-level lineages 4, 7, and 9 and unclassified DsrAB sequences from the *Firmicutes* group and the *Nitrospira* supercluster (Figure 2 B). At NGI stations, several phyla/classes, i.e., *Acidobacteria, Bacteroidetes, Deltaproteobacteria, Dependentiae* (TM6), *Gammaproteobacteria, Omnitrophica* (OP3), *Planctomycetes*, and *Woesarchaeota* (DHVEG-6) decreased in relative abundance with depth, particularly at station 3 (Figure 2 A). The phyla *Atribacter* (JS1), *Aerophobetes* (BHI80–139), *Aminicenantes* (OP8), *Alphaproteobacteria, Betaproteobacteria*, and *Chloroflexi* increased in relative 16S rRNA gene abundances with depth at both NGI stations (Figure 2 A). The relative abundances of the following *dsrAB*-containing groups decreased with depth at NGI stations: *Desulfobacteraceae*, *Syntrophobacteraceae*, the uncultured family-level lineages 7 and 9, and uncultured bacteria within the Environmental supercluster 1 (Figure 2 B). In contrast, representatives of the uncultured family-level lineage 3, as well as uncultured bacteria within the *Deltaproteobacteria* supercluster and the *Firmicutes* group, increased in relative abundances with depth at the NGI stations.

**Table 2.** Mantel correlations between 16S rRNA gene and *dsrB* community compositions and physicochemical parameters. Only significant values (p < 0.05) are shown in the table. Parameters with the strongest effect on the 16S rRNA gene or *dsrB* community compositions are indicated in bold.

|  | 16S rRNA gene |  | *dsrB* |  |
| --- | --- | --- | --- | --- |
|  | Mantel statistic | p-value | Mantel statistic | p-value |
| Sediment age [actual years] | **0.380** | 0.003 | **0.538** | 0.002 |
| Sulfate [mmol L<sup>-1</sup>] | 0.179 | 0.016 | 0.390 | 0.002 |
| DIC [mmol L<sup>-1</sup>] | 0.108 | 0.047 | 0.207 | 0.010 |
| H<sub>2</sub>S [μmol L<sup>-1</sup>] | 0.227 | 0.003 | 0.418 | 0.002 |
| Fe(II) [μmol L<sup>-1</sup>] | 0.262 | 0.003 | - | - |
| Density [g cc<sup>-1</sup>] | 0.234 | 0.003 | 0.349 | 0.002 |
| Porosity | 0.213 | 0.003 | 0.364 | 0.002 |
| TOC [μmol g dw<sup>-1</sup>] | 0.228 | 0.005 | 0.329 | 0.002 |
| C/N ratio | **0.471** | 0.003 | **0.478** | 0.002 |
| SRR [nmol Sulfate cm<sup>-3</sup> d<sup>-1</sup>] | 0.260 | 0.003 | - | - |
| Methane [μmol L<sup>-1</sup>] | - | - | 0.310 | 0.002 |

DIC, dissolved inorganic carbon; TOC, total organic carbon; SRR, sulfate reduction rate

**Figure 2.**
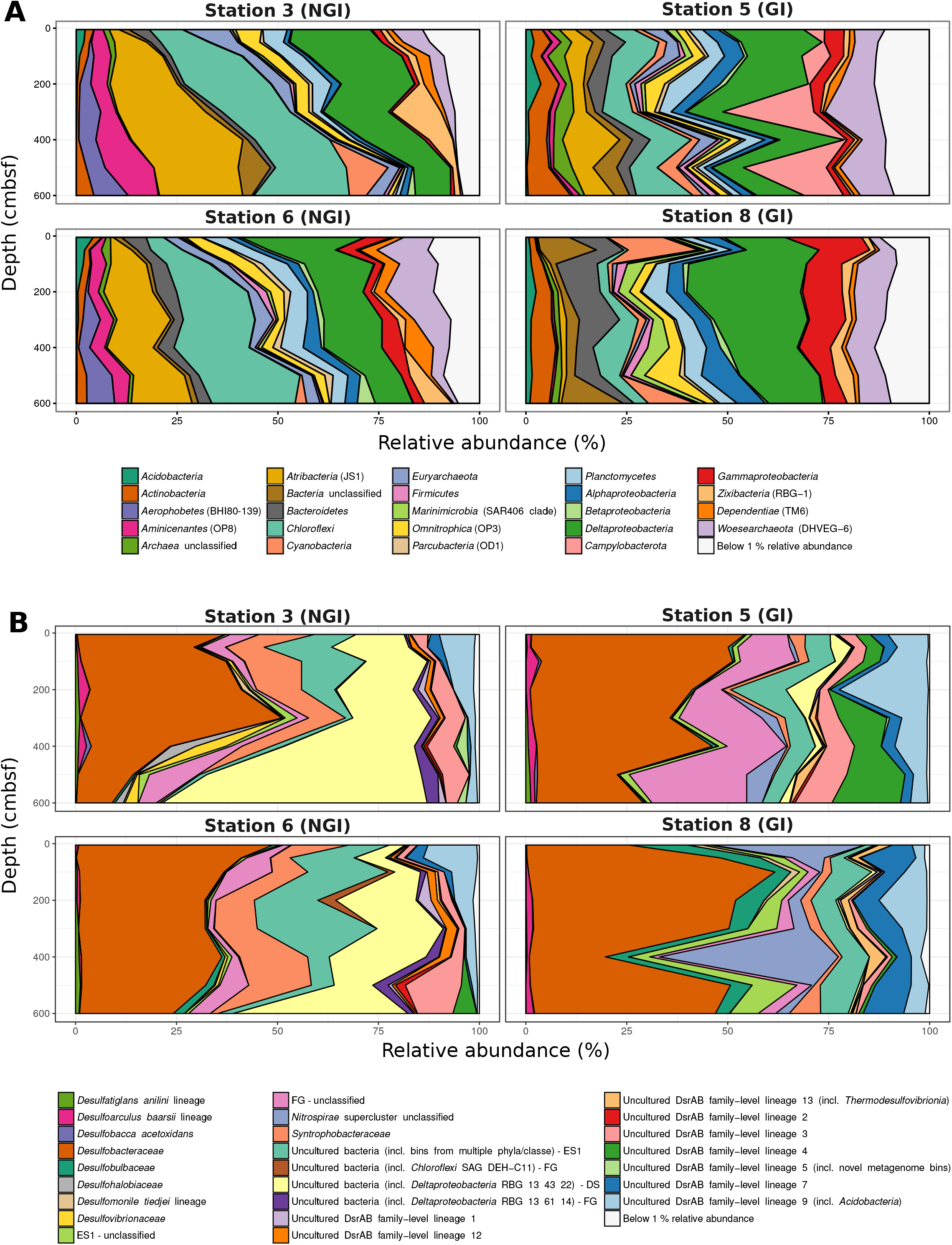
Microbial community composition in non-glacier-influenced (NGI) and glacier-influenced (GI) sediment cores. Changes in 16S rRNA phylum/class (A) and DsrAB-family (B) relative abundances with sediment depth are shown. Only phyla/classes and DsrAB-families with a relative abundance greater than 1% are shown. DS, *Deltaproteobacteria* supercluster. ES1, Environmental supercluster 1. FG, *Firmicutes* group. NS, *Nitrospira* supercluster.

### OTUs that were positively correlated to SRR have high inter-species connectivity in young sediments

Correlation network analyses of 16S rRNA- and *dsrB*-OTUs revealed two nearly separated 16S rRNA-OTU clusters and two completely separated *dsrB*-OTU clusters, which were structured along a sediment age gradient (Figure 3 A and B). One cluster included OTUs that were most abundant in old NGI sediments and the other one included OTUs that were most abundant in young GI and NGI sediments. These separate network clusters in ‘young’ and ‘old’ sediments largely overlapped with regions of particularly high inter-species OTU correlations. These ‘young’ and ‘old’ sediment OTU clusters were separated at a sediment age of around 300-400 years (Figure 3 C and D), which corresponds to a sediment depth of 30-40 and 20-30 cmbsf at NGI stations 3 and 6, respectively.

**Figure 3.**
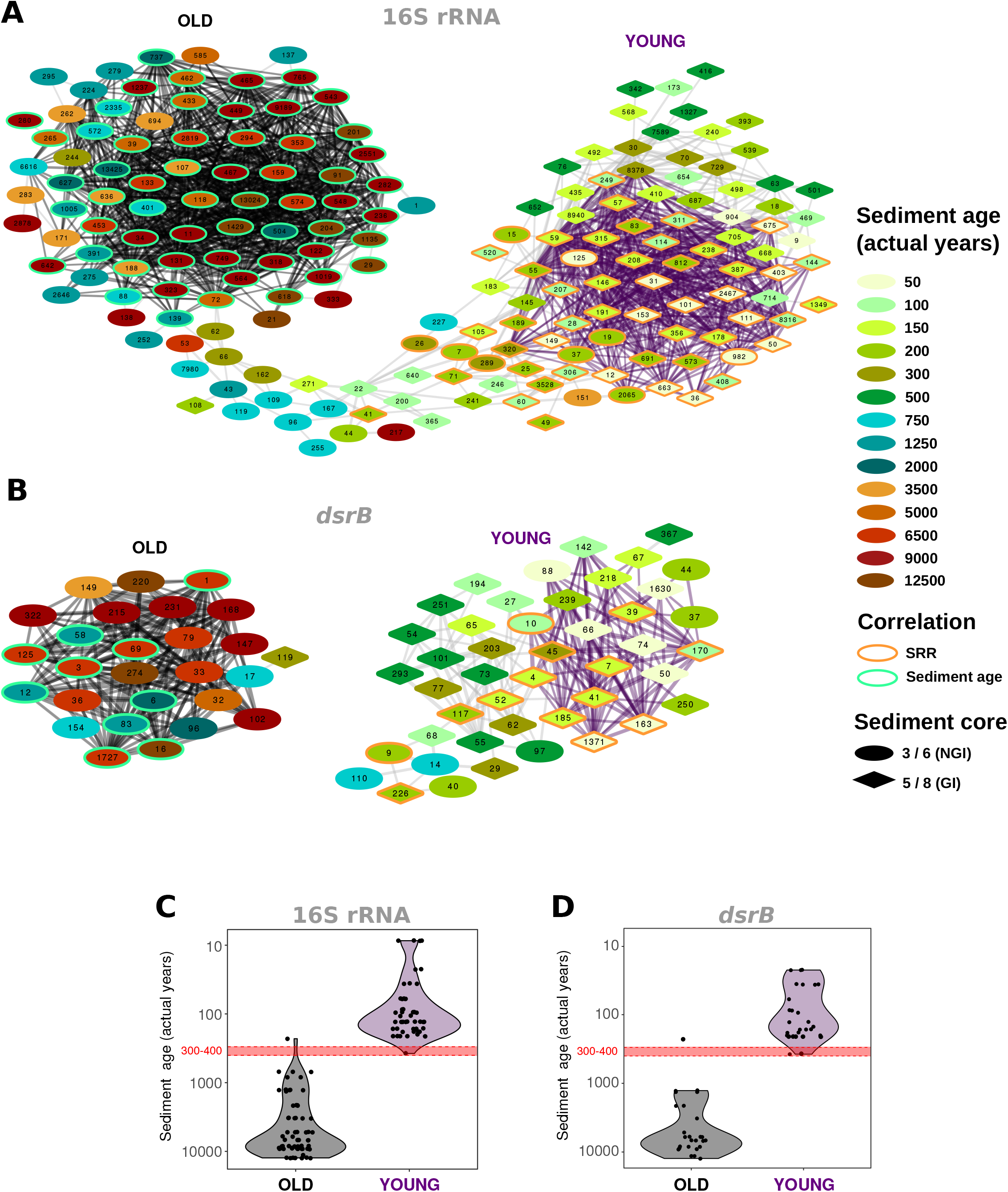
Co-occurrence of abundant OTUs across non-glacier-influenced (NGI) and glacier-influenced (GI) sediment cores. Inter-species correlations are indicated for 16S rRNA gene (A) and *dsrB* (B) OTUs. Only edges with p ≤ 0.01 and R^2^ ≥ 0.5 are shown. OTUs are colored and shaped according to the approximated sediment age and sampling station at which they were found at the highest relative abundance, respectively. The orange and green border color of OTUs indicates significant correlations to sulfate reduction rates and sediment age, respectively. OTUs that are connected by black and purple edges formed significant community clusters in old and young sediments, respectively. C and D, Age of the sediment layer at which individual OTUs from (A) and (B) were found at the highest relative abundance across all samples and stations. Each dot represents an OTU. Black and purple background colors indicate the affiliation to significant community clusters determined for the inter-species correlation networks, respectively.

The OTUs in the 16S rRNA- and *dsrB*-OTU networks were additionally subjected to correlation analyses with environmental parameters (Supplementary Figures S3 and S4). The majority of OTUs that constituted the ‘young’ and ‘old’ sediment clusters were positively correlated with SRR and sediment age, respectively (Figure 3 A and B). Most OTUs that correlated positively with SRR also correlated negatively with sediment age and vice versa (Supplementary Figures S3 and S4). OTUs that positively correlated to SRR showed distinct distributions in relative abundances, i.e., highest abundances in the surface sediments of NGI stations and mostly ubiquitous distributions throughout the whole core at GI stations (Figure 4). Many of these 16S rRNA-OTUs (*n*=10) and most of the *dsrB*-OTUs belonged to the family *Desulfobacteraceae* (Supplementary Figures S5 and S6). Other SRR-correlated 16S rRNA-OTUs belonged to the families *Desulfobulbaceae*, *Desulfarculaceae*, and *Syntrophobacteraceae* and to the phyla/classes, *Acidobacteria*, *Actinobacteria*, *Alphaproteobacteria*, *Bacteroidetes*, *Ignavibacteria, Gammaproteobacteria, Planctomycetes* and *Woesarchaeota* (Supplementary Figure S5). Besides the prevalence of *Desulfobacteraceae*, *dsrB*-OTUs positively correlated to SRR were affiliated with the uncultured family-level lineages 7 and 9 (Supplementary Figure S6). The *dsrB*-OTUs 40 and 41 were also affiliated with the Environmental Supercluster 1. OTU 40 belongs to a sequence cluster of uncultured bacteria, and is related to the metagenome-derived genome RBG_13_60_13 (accession number GCA_001796685.1) of a *Chloroflexi* bacterium (Supplementary Figure S6).

**Figure 4.**
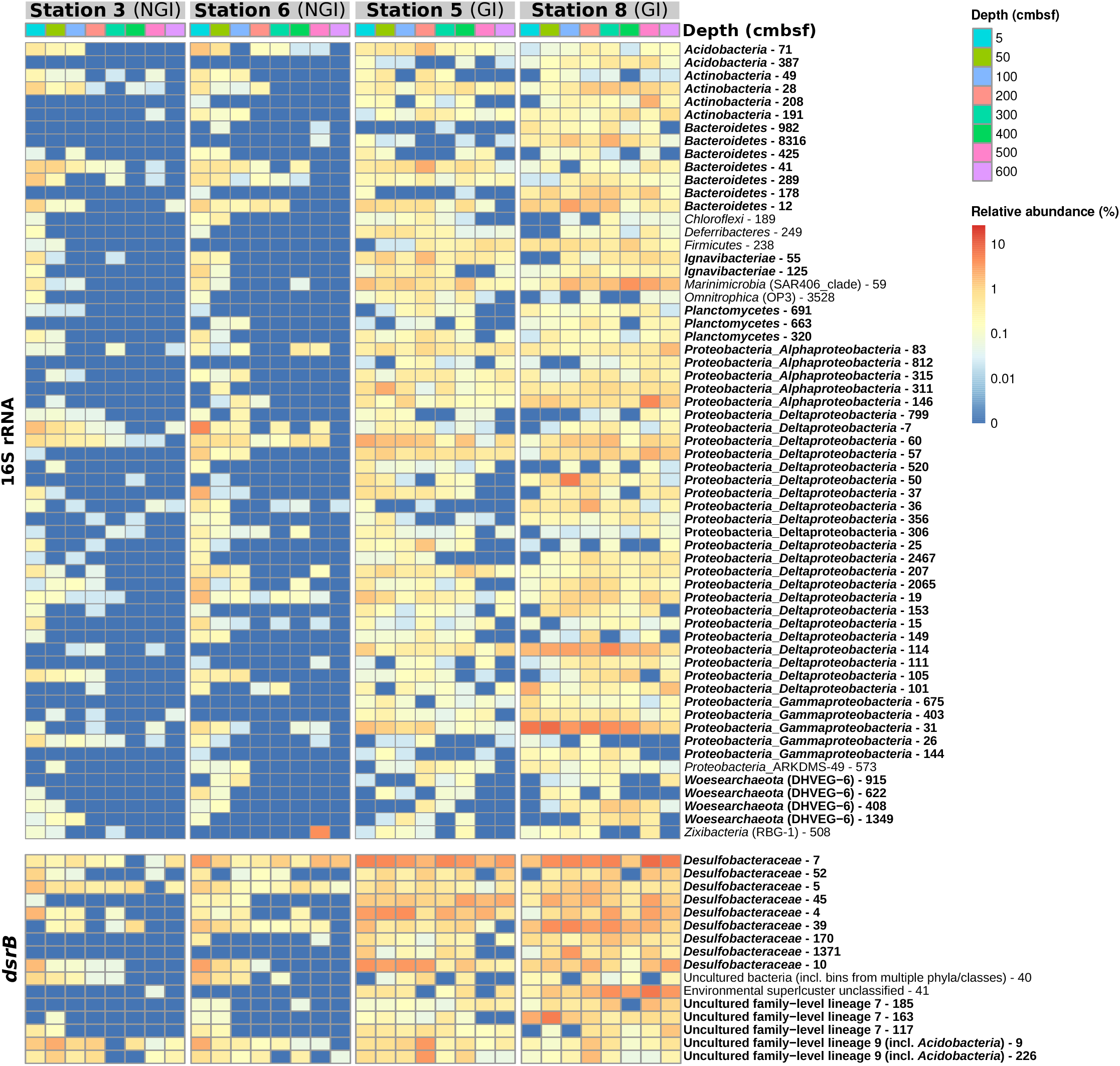
Relative abundances of 16S rRNA- and *dsrB*-OTUs with significant correlations to sulfate reduction rates in non-glacier-influenced (NGI) and glacier-influenced (GI) sediment cores. The column annotation indicates the sampling depth in centimeters below seafloor (cmbsf). The color range from blue to red indicates the relative abundance of OTUs. Phyla/classes and DsrAB-families that were represented by more than one OTU are indicated in bold.

OTUs positively correlated with sediment age were affiliated with diverse taxa (Figure 5). The phyla/classes that were represented by at least two age-correlated 16S rRNA-OTUs were *Aerophobetes* (BHI80-13), *Alphaproteobacteria, Aminicenantes* (OP8), *Atribacter* (JS1), *Chloroflexi*, *Deltaproteobacteria*, *Euryarchaeota (Marine Benthic Group D), Omnitrophica* (OP3) and *Planctomycetes* (Supplementary Figure S5). Three *dsrB*-OTUs that positively correlated with sediment age were affiliated to the family *Desulfobacteraceae* (Supplementary Figure S6). The *dsrB*-OTU 16 belonged to a group of uncultured bacteria in the *Deltaproteobacteria* supercluster, OTU 58 could only be assigned to the *Firmicutes* group, and OTU 1 was affiliated with the family *Syntrophobacteraceae*.

**Figure 5.**
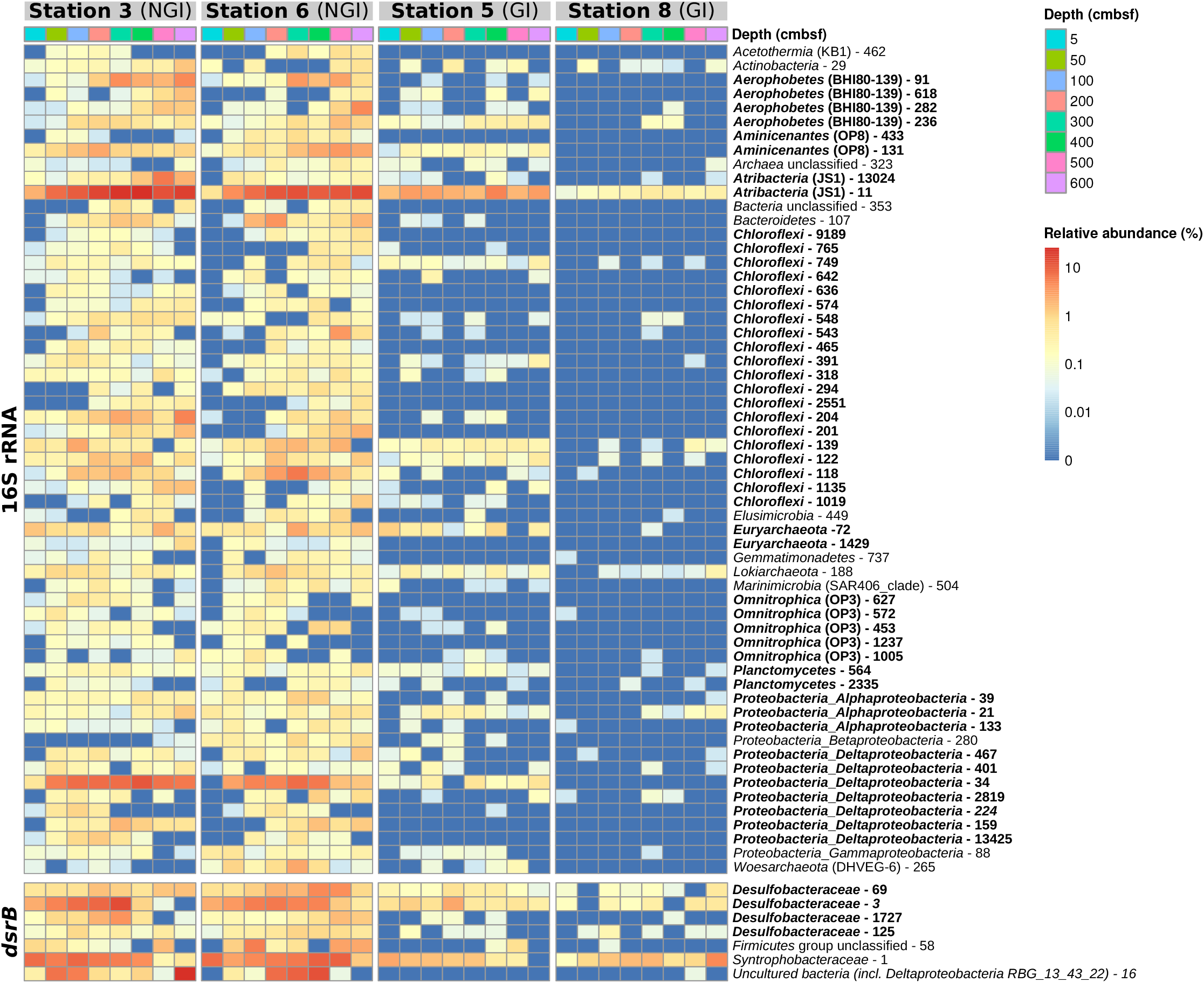
Relative abundances of 16S rRNA- and *dsrB*-OTUs with significant correlations to sediment age in non-glacier-influenced (NGI) and glacier-influenced (GI) sediment cores. The column annotation indicates the sampling depth in centimeters below seafloor (cmbsf). The color range from blue to red indicates the relative abundance of OTUs. Phyla/classes and DsrAB-families that were represented by more than one OTU are indicated in bold.

## Discussion

### Glacier-runoff affects age-depth relationships and microbial community assembly in marine sediments

Rates and amounts of glacial inputs into sedimentary environments of fjords have a major impact on their biogeochemistry (Glombitza et al., 2015). Here, we have compared microbial community structure between NGI and GI sediments, and highlighted the environmental factors that underlie the observed differences in community assembly with sediment depth. Sulfate is the key terminal electron acceptor in marine sediments of the Godthåbsfjord (Sørensen et al., 2015), but the depth distributions of SRR and microbial community structures differed between NGI and GI sediments.

In the NGI sediments, steep gradients of SRR (Figure 1) indicated that most of the labile organic matter deposited from marine primary production was mineralized near the seafloor surface as typically observed for marine shelf sediments (Flury et al., 2016). Microbial communities in the NGI sediments became less diverse with depth and increasingly distinct from the surface communities (Supplementary Figure S2). Such shifts in community composition with depth can be attributed to the progressing geochemical stratification of the sediment and decreasing flux of energy with increasing sediment age (Petro et al., 2017).

In contrast, the two GI sediment cores were characterized by higher sedimentation rates, low porosity (station 8), young ages, low TOC concentrations, low TN, and high C/N ratios (Figure 1); the latter being attributed to influx of terrestrial organic matter (Goñi et al., 2013; Meyers, 1994; Wehrmann et al., 2014). Terrestrial organic matter of particulate phase transported by the glaciers is mostly old, diagenetically altered, and likely unavailable for microbial degradation (Wehrmann et al., 2014). Therefore, the strong impact of C/N ratio differences on the microbial community might only reflect the high rate of sedimentation. The on average higher SRR throughout the GI sediment cores were possibly sustained by low amounts of reactive organic matter that was deposited from algal blooms in nutrient-rich waters of glaciated fjords (Bourgeois et al., 2016). Glacial runoff also contains considerable amounts of iron and manganese (Bhatia et al., 2013; Wehrmann et al., 2014). Accumulation of dissolved Fe^2+^ suggested that Fe(III) reduction substantially contributed to organic matter mineralization in upper GI station sediments (Wehrmann et al., 2014). Lack of sulfide accumulation with depth indicated immediate re-oxidation and/or scavenging of sulfide produced from high sulfate reduction activity (Figure 1) (Wehrmann et al., 2017). In agreement with previous studies (Bourgeois et al., 2016; Buongiorno et al., 2019; Park et al., 2011), differences in organic matter availability and electron acceptor concentrations are suggested to had a major influence on the composition of the seafloor microbial community in glaciated fjords.

### Identities and potential functional interactions of sulfur cycling-associated taxa in ‘young’ NGI and GI sediments

16S rRNA gene and *dsrB* correlation network analyses both revealed two main OTU interaction clusters (Figure 3). In one cluster most OTUs were positively correlated to SRR but not sediment age, while in the other cluster most OTUs were positively correlated to sediment age but not SRR. The relative abundances of 16S rRNA- and *dsrB*-OTUs that positively correlated with SRR were highest in ‘young’ GI and NGI sediments with active sulfur cycling (up to about 400 years of age). The majority of these OTUs was affiliated with the family *Desulfobacteraceae*, as well as other *bona fide* deltaproteobacterial SRM taxa from marine sediments (Wasmund et al., 2017). In addition, several SRR-correlated *dsrB*-OTUs were affiliated with uncultured DsrAB lineages. Some of these lineages contain metagenome-derived sequences of uncultivated bacteria from the phyla *Acidobacteria* (DsrAB family-level lineage 9), *Planctomycetes* or *Chloroflexi* (Anantharaman et al., 2017; Wasmund et al., 2016). Interestingly, several 16S rRNA-OTUs that positively correlated with SRR were also affiliated with these phyla, supporting their putative involvement in sulfite/sulfate reduction or sulfur disproportionation in the Godthåbsfjord sediments.

Despite high rates of sulfate reduction, sulfide did not accumulate in the GI sediments (Figure 1) likely due to its reaction with metals resulting in its oxidation and or precipitation (Wehrmann et al., 2014). Several 16S rRNA-OTUs that correlated positively with SRR were affiliated with taxa containing sulfur-oxidizing microorganisms such as the candidate genus PHOS-HE36 (phylum *Ignavibacteriae*) (Koenig et al., 2005), the *Woeseiaceae*/JTB255 sediment group (*Gammaproteobacteria*) (Dyksma et al., 2016; Mußmann et al., 2017), and the *Rhodobacteriaceae* (*Alphaproteobacteria*) (Lenk et al., 2012; Thrash et al., 2017) (Supplementary Figure S5). In addition, OTU 30 affiliated with the sulfur-oxidizing genus *Sulfurovum* was solely responsible for the high relative abundance of *Campylobacterota* at GI station 5. We hypothesize that high SRR and chemical oxidation of sulfide by metals may support significant populations of sulfur-oxidizing or sulfur-disproportionating taxa in deep GI sediments, although future work would be required to substantiate this hypothesis, e.g., detection of mRNA transcripts for sulfur-dissimilating enzymes.

We also identified numerous SRR-correlated OTUs that were affiliated with taxa that are not known to have sulfur-based energy metabolisms. While these OTUs could indeed represent unknown sulfur-cycling microorganisms, they may also be degraders of organic matter that fuel sulfate reduction with fermentation products. For instance, some of these 16S rRNA-OTUs were affiliated with BD2-2 (phylum *Bacteroidetes*), *Phycisphaera* (phylum *Planctomycetes*) or OM1 (phylum *Actinobacteria*). Representatives of these phyla hydrolyze and ferment organic polymers in marine sediments and consequently might have trophic associations with SRM (Baker et al., 2015; Schauer et al., 2011; Trembath-Reichert et al., 2016; Webster et al., 2011). These associations may therefore explain their cooccurrences with SRM detected here.

### Assembly of the deep subsurface microbial biosphere in NGI sediments

OTU correlation analysis showed that OTU clusters and thus the microbial communities of young and old sediment zones in NGI sediments became ‘disconnected’ at a sediment age of about 300-400 years (Figure 3), corresponding to a sediment depth of approximately 30 cmbsf at NGI stations. This is in line with observations that a considerable shift in microbial community structures occur in marine sediments below the zone of bioturbation, which was suggested to be the main site of assembly of the subsurface community (Jochum et al., 2017; Starnawski et al., 2017). Most OTUs that positively correlated with sediment ages in NGI sediments were affiliated with lineages known to harbor members that selectively persist from the surface into deep subsurface sediments, e.g., *Chloroflexi, Aerophobetes, Atribacter* (JS1), *Aminicenantes* (OP8), *Alphaproteobacteria* and *Deltaproteobacteria* (Supplementary Figure S5) (Orcutt et al., 2011; Wang et al., 2016). Members of these taxa, such as the genus *Desulfatiglans* (Jochum et al., 2018), the deltaproteobacterial candidate lineage SEEP-SRB1 (Schreiber et al., 2010) or the euryarchaeal Marine Benthic Group D (Kaster et al., 2014; Lloyd et al., 2013; Nobu et al., 2016; Robbins et al., 2016; Wang et al., 2016b; Wasmund et al., 2014) are postulated to have traits such as fermentation, sulfate reduction or acetogenesis to support the maintenance of basic cellular functions even under extreme energy-limited conditions in most subsurface sediment environments (Petro et al., 2017).

## Conclusions

Coastal marine ecosystems in arctic and sub-arctic oceans are poised to be increasingly impacted by melting of glaciers caused by climate change. In this comparative study, we found that discharge from marine-terminating glaciers had a strong control over the depth-dependent microbial community assembly in sediments of a sub-arctic fjord. Increasing differences in the benthic community composition between GI and NGI sites with depth were largely explained by sediment age. High sedimentation rates at GI stations enabled a complex community of sulfur-cycling-associated microorganisms, including both putative SRM and sulfide oxidizers, to continuously thrive at high relative abundances from the surface deep into the subsurface. Similar communities of sulfur-cycling-associated microorganisms were also present in surface sediments at NGI stations. However, with increasing depth the surface communities were largely replaced by microorganisms that positively correlated with sediment age. Lower sedimentation rates at the NGI sites thus resulted in slow burial and highly selective survival of microorganisms adapted to the energy-limited subsurface (Petro et al., 2017). In summary, our results suggest that increased glacier runoff and the associated high sedimentation rates allow processes that are typically predominant in surface sediments such as sulfide oxidation and associated community members to be rapidly buried and maintained at high abundances in deep subsurface sediments.

## Author Contributions

ClP, HR, KUK, and AL designed the research. ClP generated and analyzed the sequencing data. MSS, HR, and ChP collected the sediment cores. MJ, HR, and KK obtained the samples and MJ performed most biogeochemical analyses. KUK performed DNA extractions. MSS, ChP, and ZAK calculated sediment ages. ClP, KW, and AL wrote the manuscript. All authors revised the manuscript.

## Acknowledgments

The authors thank the crew of the R/V Sanna and the scientific party during the 2013 sampling campaign and Britta Poulsen and Susanne Nielsen for laboratory technical assistance. We acknowledge the use of imagery from the NASA Worldview application (https://worldview.earthdata.nasa.gov), part of the NASA Earth Observing System Data and Information System (EOSDIS).

