## Supplementary Material for "Glacial runoff promotes deep burial of sulfur cycling-associated microorganisms in marine sediments"

Claus Pelikan, Marion Jaussi, Kenneth Wasmund, Marit-Solveig Seidenkrantz, Christof Pearce, Zou Zou Anna Kuzyk, Craig W. Herbold, Hans Røy, Kasper Urup Kjeldsen, and Alexander Loy

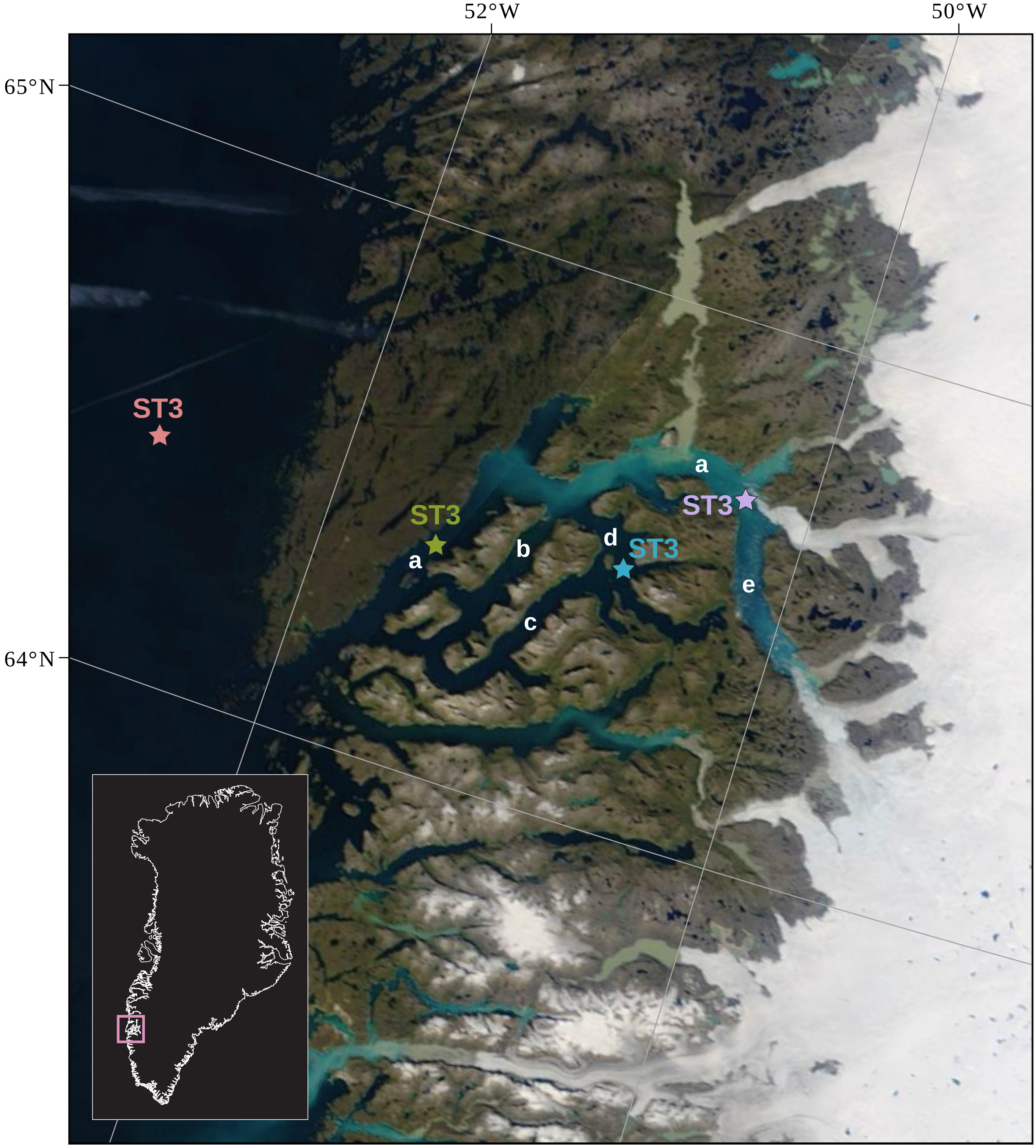


**Supplementary Figure S1. Locations of the four sampling stations in the Godthåbsfjord region.** The exact coordinates of the stations are indicated in Table 1. ST3 and ST6, Non-glacier-influenced stations 3 and 6. ST5 and ST8, Glacier-influenced stations 5 and 8. a, main fjord (Nûp Kangerdlua). b, Qôrnup Suvdlua. c, Ũmánap Suvdlua. d, Kapisigdlit Kanderdluat. e, Kangersuneq. The background map is a NASA Worldview satellite image (Terra MODIS Corrected Reflectance) of 19 August 2013.

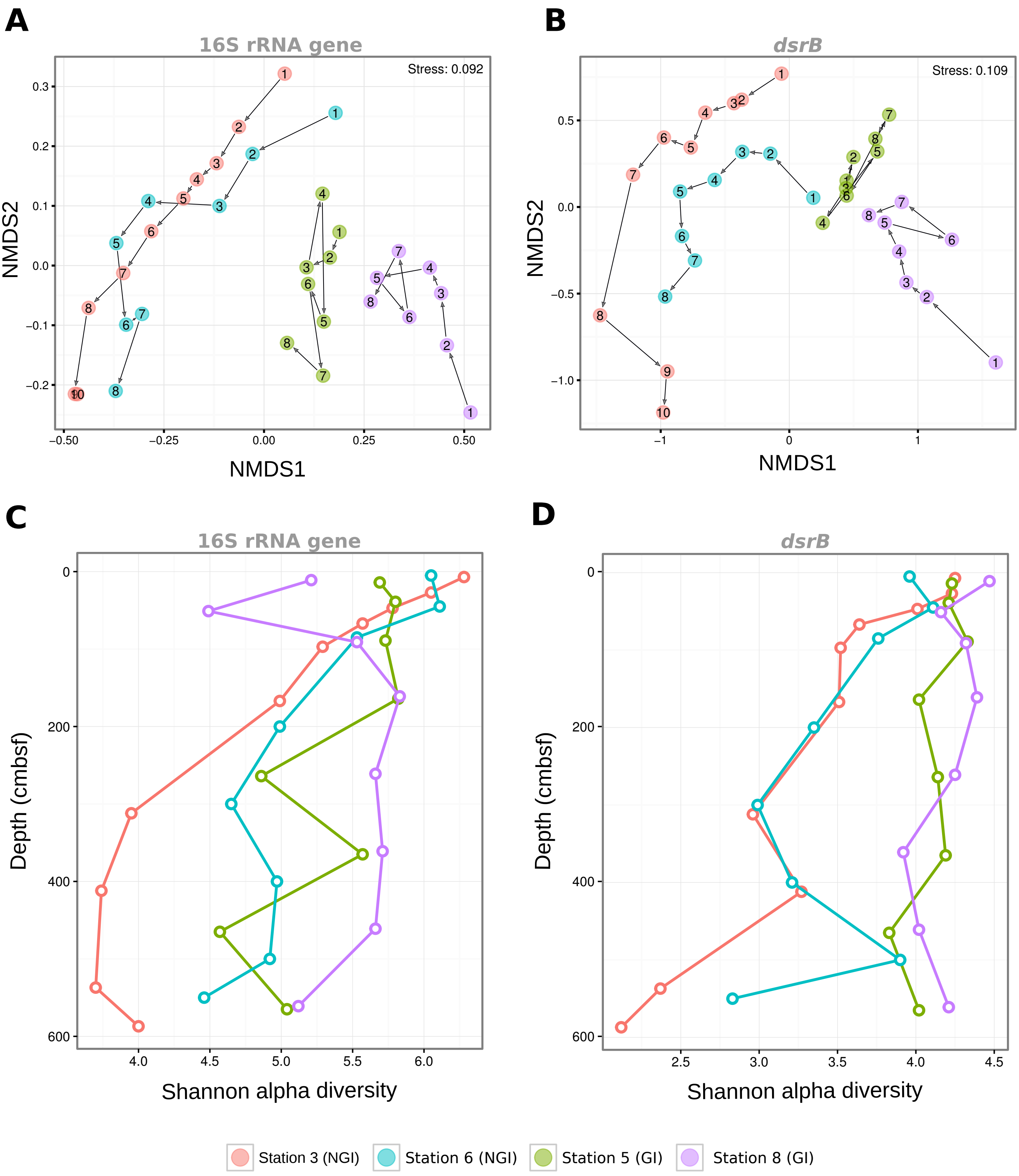


**Supplementary Figure S2. Microbial diversity across non-glacier-influenced (NGI) and glacier-influenced (GI) sediment cores and sediment depth.** A and B, Nonmetric multidimensional scaling ordinations (NMDS) of Bray-Curtis distances between 16S rRNA gene and *dsrB* communities. C and D, Shannon alpha diversity indices of 16S rRNA gene and *dsrB* communities. Colours indicate the sampling station. Numbers indicate the sediment depth according to Supplementary Table 1.

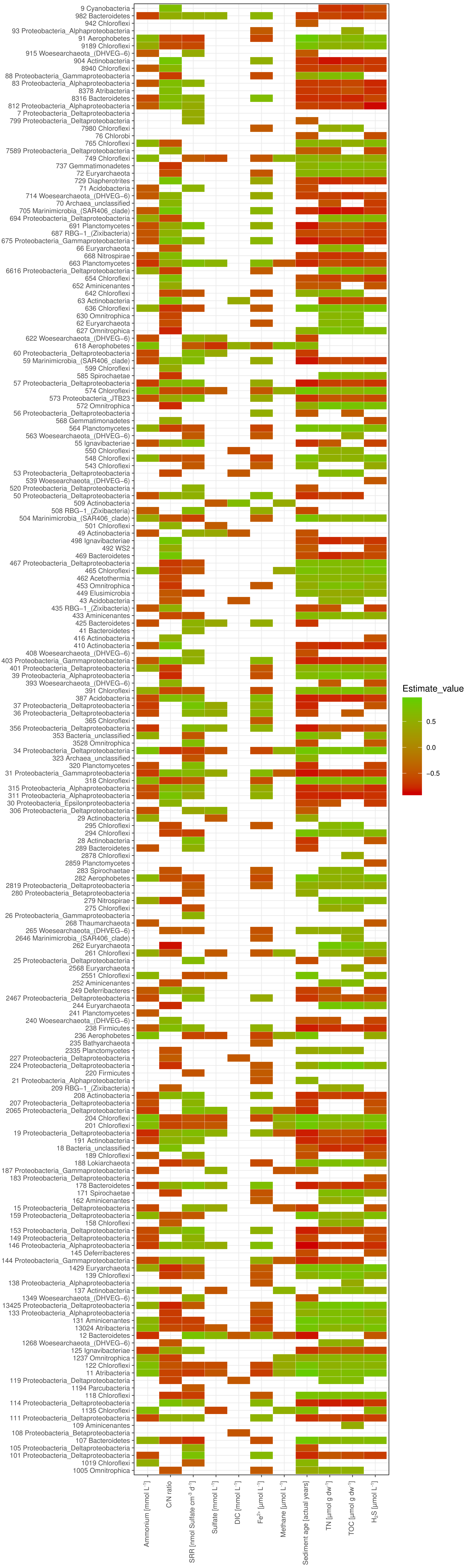


**Supplementary Figure S3. Spearman correlations between relative abundances of 16S rRNA-OTUs and physicochemical parameters.** Indicated are OTUs with significant ($p \leq 0.05$) Spearman correlations to any of the tested parameters.

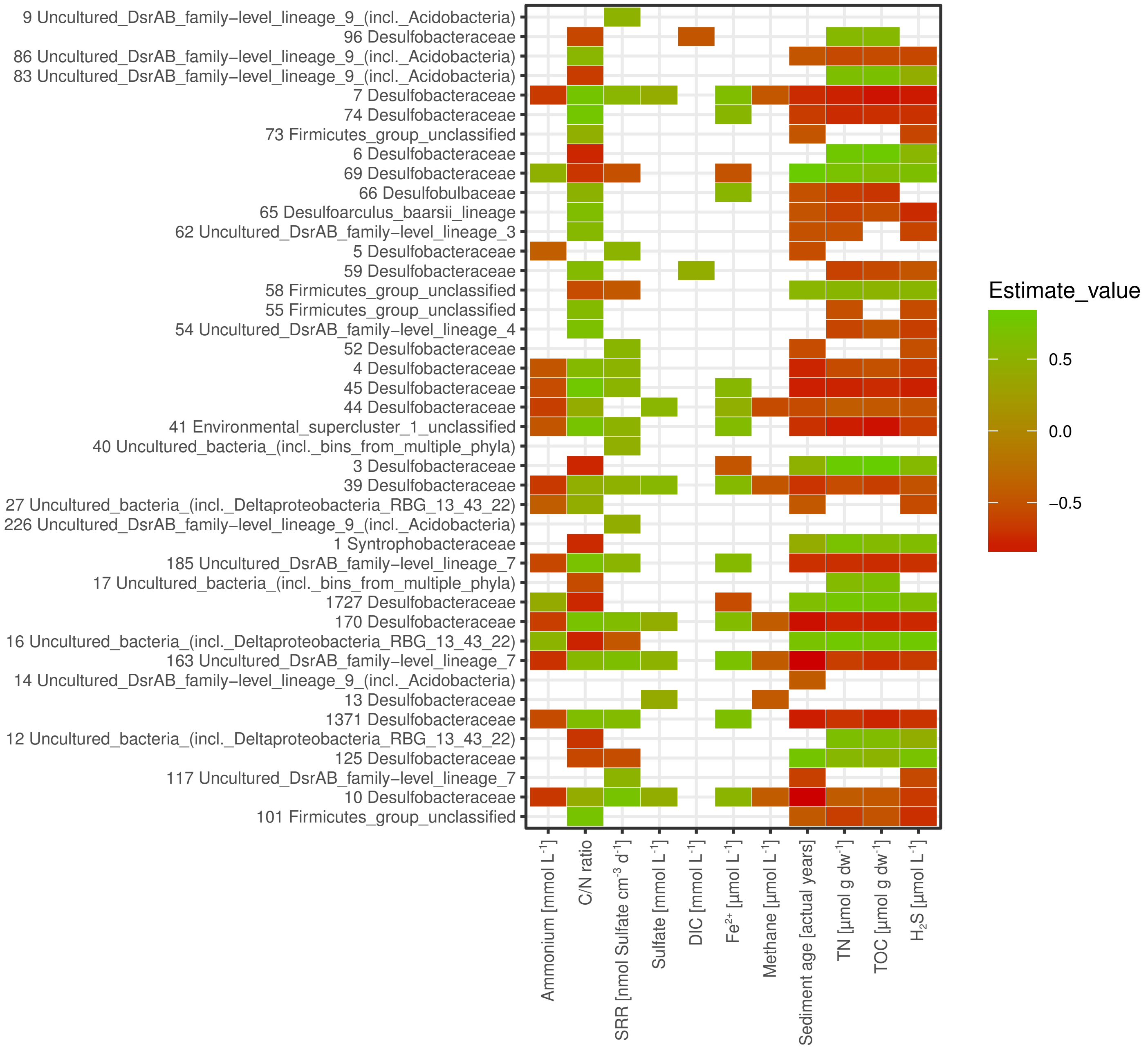


**Supplementary Figure S4. Spearman correlations between relative abundances of *dsrB*-OTUs and physicochemical parameters.** Indicated are OTUs with significant ($p \leq 0.05$) Spearman correlations to any of the tested parameters.

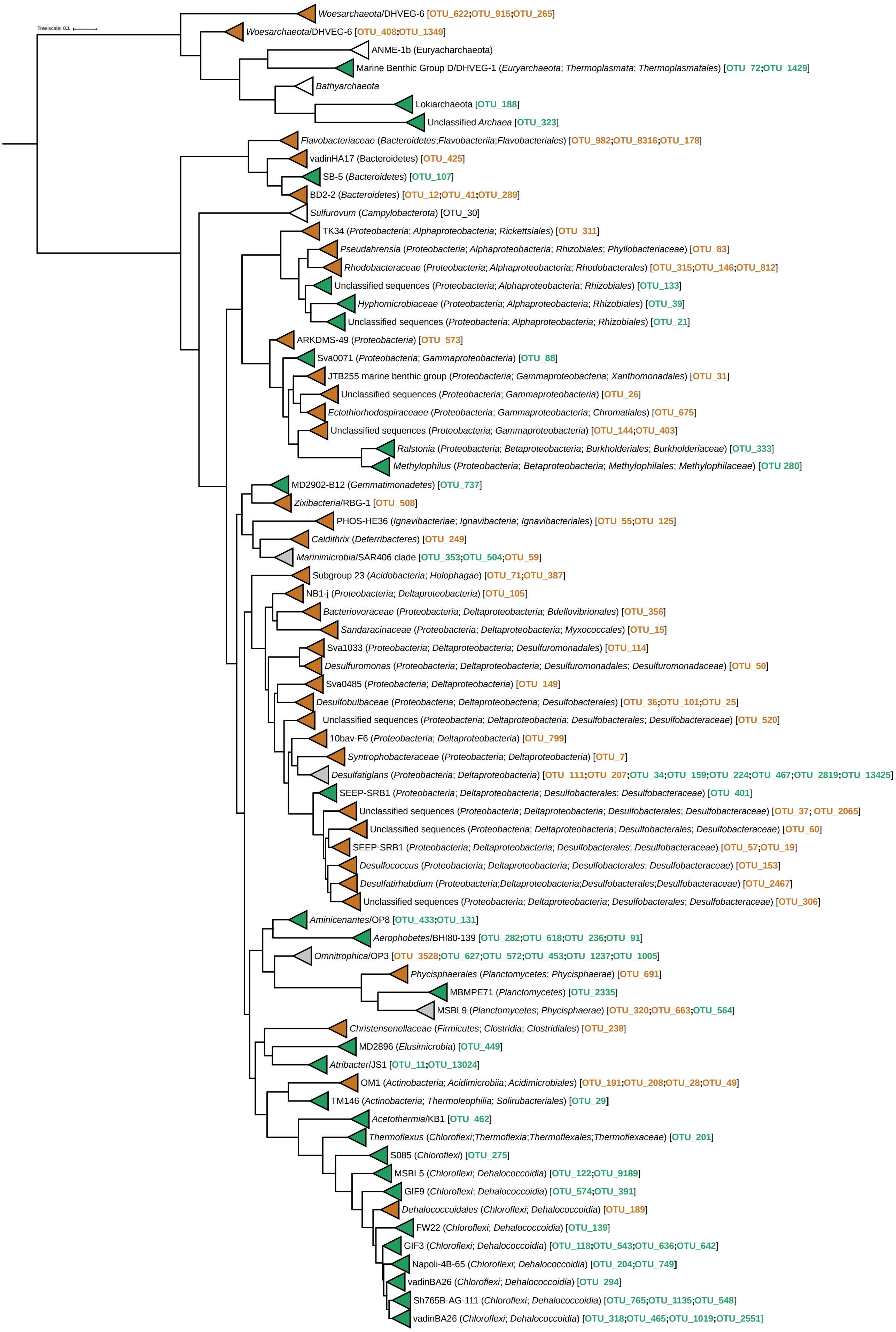


**Supplementary Figure S5. Phylogenetic placement of 16S rRNA-OTUs.** Only OTUs with significant correlations to sulfate reduction rates or sediment age are shown. The reference tree was built from sequences that were closely related to 16S rRNA-OTUs. These were extracted from the SILVA database v.128 (Quast *et al.*, 2013), and used for tree construction with FastTree (Price *et al.*, 2010). 16S rRNA-OTUs were aligned to the SILVA database using the SINA aligner (Pruesse *et al.*, 2012), and placed into the reference tree using the EPA algorithm (Berger *et al.*, 2011) in RAxML (Stamatakis, 2014). The placement tree was visualized with iTOL (Letunic and Bork, 2007). OTUs that are colored in orange and green were significantly correlated to sulfate reduction rates (Figure 3) and sediment age (Figure 4), respectively. Clades that contained only OTUs with significant correlations to either sulfate reduction rates or sediment age are colored accordingly. Gray clades contain OTUs with correlations to sulfate reduction rates and OTUs with correlations to sediment age.

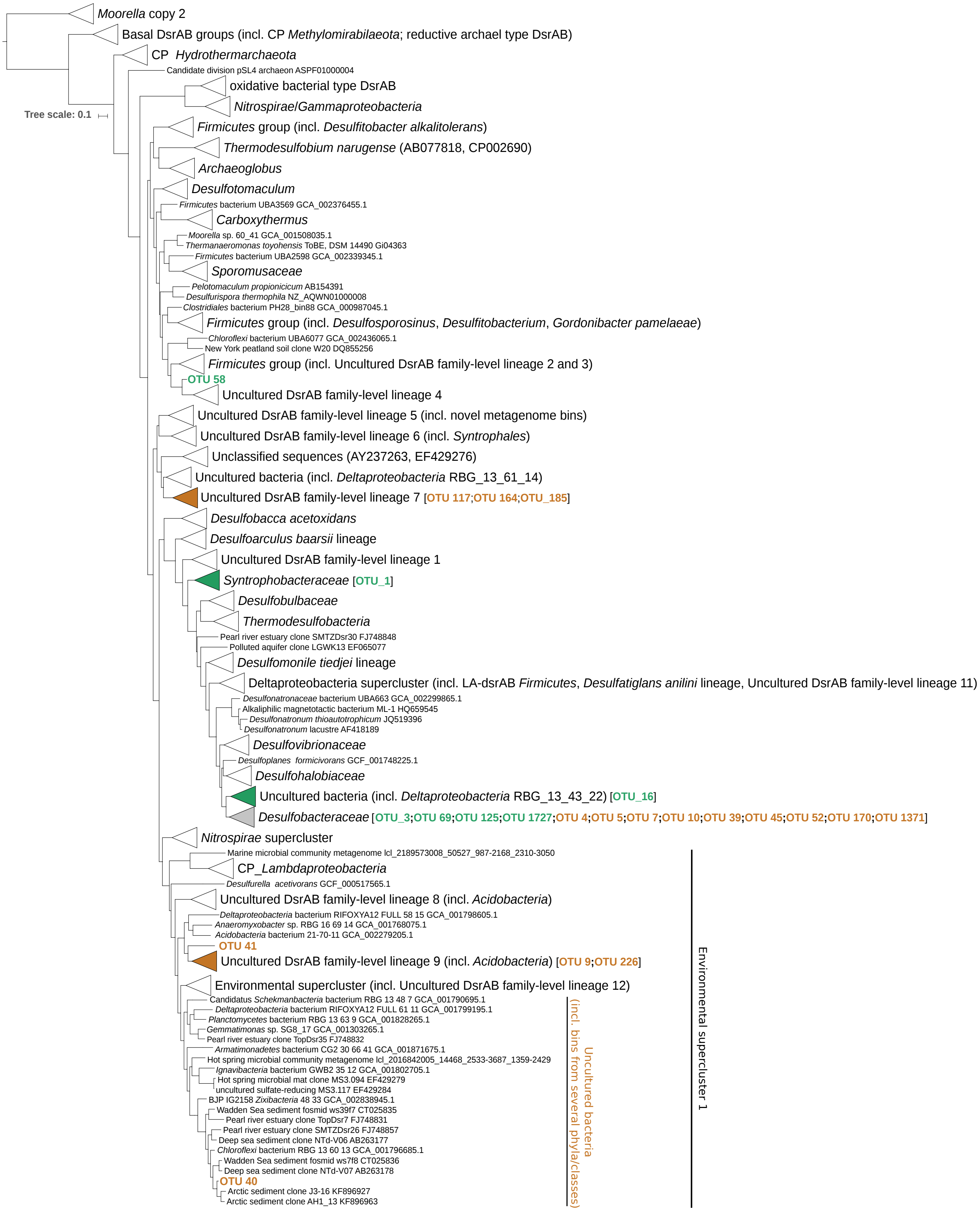


**Supplementary Figure S6. Phylogenetic placement of *dsrB*-OTUs.** Only OTUs with significant correlations to sulfate reduction rates or sediment age are shown. The reference tree was built from an extended reference alignment that contained DsrAB sequences from Müller *et al.*, 2015 and novel DsrAB sequences from recent metagenomic surveys (Anantharaman *et al.*, 2017; Parks *et al.*, 2017). The reference tree was constructed with FastTree (Price *et al.*, 2010). *dsrB*-OTUs were aligned to the extended reference alignment using MAFFT (Katoh *et al.*, 2002), and added to the reference tree using the EPA algorithm (Berger *et al.*, 2011) in RAxML (Stamatakis, 2014). The placement tree was visualized with iTOL (Letunic and Bork, 2007). OTUs that are colored in orange and green were significantly correlated to sulfate reduction rates (Figure 4) and sediment age (Figure 4), respectively. Clades that contained only OTUs with significant correlations to either sulfate reduction rates or sediment age are colored accordingly. Gray clades contain OTUs with correlations to sulfate reduction rates and OTUs with correlations to sediment age.

**Supplementary Table S1. Sample numbering and sediment depth of the four sediment cores.**

| Coring station | Sample numbering | Sediment depth (cmbsf) |
|---|---|---|
| Station 3 | 1 | 25 |
| Station 3 | 2 | 45 |
| Station 3 | 3 | 65 |
| Station 3 | 4 | 85 |
| Station 3 | 5 | 115 |
| Station 3 | 6 | 185 |
| Station 3 | 7 | 330 |
| Station 3 | 8 | 430 |
| Station 3 | 9 | 555 |
| Station 3 | 10 | 605 |
| Station 5 | 1 | 28 |
| Station 5 | 2 | 53 |
| Station 5 | 3 | 103 |
| Station 5 | 4 | 178 |
| Station 5 | 5 | 278 |
| Station 5 | 6 | 379 |
| Station 5 | 7 | 479 |
| Station 5 | 8 | 579 |
| Station 6 | 1 | 15 |
| Station 6 | 2 | 55 |
| Station 6 | 3 | 95 |
| Station 6 | 4 | 210 |
| Station 6 | 5 | 310 |
| Station 6 | 6 | 410 |
| Station 6 | 7 | 510 |
| Station 6 | 8 | 560 |
| Station 8 | 1 | 25 |
| Station 8 | 2 | 65 |
| Station 8 | 3 | 105 |
| Station 8 | 4 | 175 |
| Station 8 | 5 | 275 |
| Station 8 | 6 | 375 |
| Station 8 | 7 | 475 |
| Station 8 | 8 | 575 |
